# Identification of visible and near-infrared signature peaks for arboviruses and *Plasmodium*

**DOI:** 10.1101/2023.10.26.564151

**Authors:** Brendon Goh, Ricardo J. Soares Magalhães, Silvia Ciocchetta, Wenjun Liu, Maggy T. Sikulu-Lord

## Abstract

Arbovirus and malaria infections affect more than half of the world’s population causing major financial and physical burden. Current diagnostic tools such as microscopy, molecular and serological techniques are technically demanding, costly, or time consuming. Near-infrared spectroscopy has recently been demonstrated as a potential diagnostic tool for malaria and arbovirus and as a screening tool for disease vectors. However, pathogen specific infrared peaks that allow detection of these infections are yet to be described. In this study, we identified unique NIRS peaks from existing laboratory strains of four major arboviruses including Barmah Forest virus (BFV), Dengue virus (DENV), Ross River virus (RRV), Sindbis virus (SINV) and *Plasmodium falciparum*. Secondly, to determine the diagnostic ability of these peaks, we developed machine learning algorithms using Artificial Neural network (ANN) to differentiate arboviruses from media in which they are grown. Signature peaks for BFV were identified within the visible region at 410, 430, 562 and 588nm and the NIR region at, 946, 958, 1130, 1154 and 1780 nm. DENV related peaks were seen at 410nm within the visible region and 1130 nm within the NIR region. Signature peaks for RRV were observed within the visible region at 410 and 430 nm and within the NIR region at 1130 and 1780 nm, while SINV had a prominent peak at 410 nm within the visible region. Peaks at 514, 528, 547, 561, 582, and 595nm and peaks at 1388, 1432, 1681, 1700, 1721, 1882, 1905, 2245, 2278, 2300 nm were unique for *P. falciparum*. NIRS predictive sensitivity defined as the ability to predict an arbovirus as an infection was 90% (n = 20) for BFV, 100% (n =10) for RRV and 97.5% (n= 40) for DENV, while infection specificity defined as the ability to predict media as not-infected was 100% (n= 10). Our findings indicate that spectral signatures obtained by NIRS are potential biomarkers for diagnosis arboviruses and malaria.

**Author summary:** More than half of the world is at risk of contracting either malaria or arboviral infections. In resource limited settings, timely detection of these infections and the ability to screen thousands of people in a day using affordable tools is key to preventing their spread and unprecedented outbreaks. This emphasizes the need to develop portable, rapid, and easy to use diagnostic and surveillance tools. The near-infrared spectroscopy technique has recently been shown to be a potential tool for the diagnosis and surveillance of both malaria and arboviral infections. However, signature spectral biomarkers for these infections remain to be described. This study has identified several spectral biomarkers for DEN, RRV, BVF, SIN arboviruses and *P. falciparum parasites*. These biomarkers will assist the future assessment of this technique as a diagnostic and or surveillance tool for these infections in the field.

## Introduction

Arboviruses persist in nature through a life cycle involving a vertebrate host, an organism that carries the virus and an infected arthropod, usually mosquitos or ticks [1]. Arboviral infections have been on the rise due to increased geographical distribution and abundance of arthropod vectors mainly because of concurring factors such as climate change, migration and urbanization [2]. For instance, DENV infections is now considered the most common vector-borne virus globally affecting more than half of the world’s population [3].

Australia is home to a number of arboviral infections of public health significance and the most prevalent arboviral disease is that caused by RRV. RRV is endemic to mainland Australia and Tasmania, Cook Islands, Fiji, Samoa, New Caledonia, New Guinea and several other islands in the South Pacific region [4]. In 2015, Australia reported its largest ever recorded outbreak of 9,544 RRV cases [5]. BFV is the second most prevalent arbovirus and is the cause of epidemic polyarthritis throughout mainland Australia and Papua New Guinea [6]. The most recent estimates of BFV reported between 1995 and 2008 in Australia is 15,592 with Queensland state alone recording 8,050 cases [7]. In a more recent study, BFV was recovered from trapped mosquitoes in military training areas in Queensland [8]. SINV infections have also been reported in Australia however most SINV infections have recently been reported in Finland. In 2021 alone, 566 SINV infections were recorded in Finland [9]. Other occasional countries that report SINV include China and South Africa [10].

Current diagnosis of arboviruses include molecular methods such as reverse transcriptase-polymerase chain reaction (RT-PCR) [11] and serological techniques such as Enzyme-linked immunosorbent assay (ELISA) [12, 13]. Molecular methods are the most accurate and sensitive. For example, RT-PCR for DENV has detection limits of 10, 100, 10 and 100 copies/reaction for DENV-1,2,3 and 4, respectively. While ELISA assay for detection of DENV-2 specific antibodies has been reported to be 90% (n=20) specific and 100% sensitive (n=20) in 40 human serum samples [14]. Despite high sensitivity, these methods are restricted to the laboratory settings, can be time consuming and costly for programmatic diagnosis and surveillance purposes.

Malaria is a mosquito-borne disease caused by the *Plasmodium* parasite which is transmitted to humans through the bite of an infected female *Anopheles* mosquito. In 2021, an estimated 247 million malaria cases and 619,000 malaria related deaths were reported by the World Health Organisation highlighting malaria as a major public health concern [15].

Traditionally, malaria is diagnosed using microscopy and Giemsa-stained blood smears [16]. However, with a limit of detection of >50 parasites/μL of blood, it requires a well-trained microscopist [17]. Rapid diagnostic tests are also common diagnostic tools for malaria because they are very easy to use and do not require qualified personnel, but their sensitivity and specificity is low in detecting low parasitaemia [18, 19]. Molecular based techniques such as Polymerase Chain Reaction, quantitative-PCR, nested PCR and ELISA have been developed for malaria [20] but they are time consuming, costly and require trained personnel.

Near-infrared spectroscopy (NIRS) is a potential novel diagnostic/surveillance tool. It involves the interaction of near-infrared light with biological samples to produce a reflectance spectrum [21]. Based on chemical and structural differences between biological samples, unique spectra are produced. The spectra are a reflection of the amount and type of chemical composition of the sample and can be used to typify those samples. NIRS can therefore be used as a biomarker for biological samples.

A recent literature review demonstrated gaps in the application of NIR spectroscopy for diagnosis of arboviruses [22]. To date, four studies have shown the ability of NIR technique to detect DENV, Chikungunya, *Wolbachia* and Zika in *Ae. aegypti* mosquitoes with accuracies above 90% [23, 24]. However, only one study has demonstrated absorbance frequencies in the visible region for arboviruses . Firdous and colleagues reported that NIR peaks at 533 and 580 nm are indicative of the presence of the DNA mixture of DENV2 and DENV3 [25]. For malaria, NIR wavelengths have been reported for the hemozoin protein [26] and the ring stage of *P. falciparum* in whole blood [27]. The application of NIRS for the detection of the *plasmodium* parasite has been reported in several studies [28–30]. These studies identified prominent peaks across the wavelength range 200-1200 nm [28], 900-1700 nm [29] and 1300-2500 nm [30].

In this study, we identified NIR signature peaks for BFV, DENV, RRV, SINV and purified *P. falciparum* NIR. These signature peaks in combination are useful as a library of biomarkers for the diagnosis of arboviruses and malaria.

## Materials and methods

### Human serum

One biological replicate of serum sample (consisting of 150 mL of pooled human serum samples) was obtained from Australian Red Cross Lifeblood using the human ethics protocol approved by the University of Queensland (ethics approval number 2020001077). Following collection from donors, all samples were routinely tested for hepatitis B and C, HTLV I/II, syphilis HIV 1/2, and ABO/Rh antigens. Serum was stored at −25 °C 3–4 days prior to the experiment.

### Arbovirus and media cell culture

Cell culture media used in this study consisted of Roswell Park Memorial Institute Medium (RPMI) 1640 Medium (Sigma Life Sciences, USA) with 10% heat-inactivated fetal bovine serum (FBS) (Thermos Fisher Scientific, USA) and 1% Penicillin-Streptomycin Glutamine solution (PSG) (Thermos Fisher Scientific, USA). All arboviruses used were passaged in 3 separate batches. Samples from each batch were used as a single biological replicate (Table 1). Virus stocks were titrated using a modification of the Enzyme-linked immunosorbent assay procedure of Broom et al. [31]. Briefly, virus stocks and samples were 10-fold serially diluted and inoculated onto monolayers of C6/36 cells grown in C6/36 cell culture media which consisted of RPMI, 1640 Medium (Sigma Life Sciences, USA) with 5% heat-inactivated FBS (Thermos Fisher Scientific, USA) and 1% PSG (Thermos Fisher Scientific, USA) and maintained at 30 °C, 5% CO2. After 7 days of incubation, cells were fixed in acetone: methanol (1:1) for 1 hr at 4 °C. Plates were air-dried and antigen was detected using a cocktail of anti-flavivirus monoclonal antibody hybridoma supernatants; 4G2 [32] 6B-6C1:3H5 [33], at a ratio of 1:1:1, followed by horseradish peroxidase (HRP-) conjugated goat anti-mouse polyclonal antibody (DAKO, Carpinteria, CA, USA) (1:2000 in PBS-Tween 20). Antibodies bound to the cell mono-layers were detected by the addition of 3,3’,5,5’-tetramethylbenzidine liquid substrate system for membranes (Sigma-Aldrich). The cell culture infectious dose 50% (CCID50) was determined from titration endpoints as described elsewhere [34] and expressed as C6/36 CCID50/mL. This experiment was repeated three times at three separate time points to create three independent biological samples.

### Dengue virus culture

DENV prototype strains DENV-1 Hawaii (1944), DENV-2 NGC (1944), DENV-3 H-87 (1956) and DENV4 H241 (1956) were used in this study. DENV was propagated in C6/36 *Ae. albopictus* cells, maintained at 28°C in RPMI and supplemented with 10% FBS and 1% PSG. Following three passages in C6/36 cells, virus stocks were concentrated using Ultracel-100k filters (Amicon, Tullagreen, Cork Ireland) [35] and frozen at -80°C until further use.

### Barmah Forest Virus and Ross River Virus culture

BFV QML and BFV WEN 1631 and RRV QML1 strain (GenBank No. GQ433354) were used in this study. The virus strains were passaged three times in Vero cells, maintained at 37°C in RPMI and supplemented with 10% FBS and 1% PSG. Following three passages in Vero cells, virus stocks were concentrated using Ultracel-100k filters (Amicon, Tullagreen, Cork Ireland) [35] and frozen at -80°C until further use. One vial of the viral stocks was thawed to determine virus titre using 50% tissue culture infectious dose (TCID50/ml) on Vero cells as described [36]. Briefly, virus stocks were 10-fold serially diluted and 100µl of diluted virus was inoculated onto monolayers of Vero cells grown in 96 well plates in cell culture media and maintained at 37°C, 5% CO2. Ninety-six hours later, cells were fixed with 3.7% formaldehyde, stained with 1% crystal violet for 1 hour, washed in tap water and dried. The TCID50 was determined from titration endpoints as described elsewhere [37] and expressed as the Vero cell TCID50/mL.

### Sindbis virus culture

SINV strain (SINV 18953) was propagated in C6/36 *Ae. albopictus* cells, maintained at 28°C in RPMI and supplemented with 10% FBS and 1% PSG. Following three passages in C6/36 cells, virus stocks were concentrated using Ultracel-100k filters (Amicon, Tullagreen, Cork Ireland) and frozen at -80°C until further use. One vial of the viral stocks was thawed to determine virus titre by 50% tissue culture infectious dose (TCID50/ml) on Vero cells as described by Sudeep et al [38]. Briefly, virus stocks were serially diluted 10-fold and 100µl of diluted virus was inoculated onto monolayers of Vero cells grown in 96 well plates in RPMI 1640 supplemented with L-glutamine, 5% FBS, 1% PSG and maintained at 30 °C, 5% CO_2_. After 4 days of incubation, cells were fixed with 3.7% formaldehyde, stained with 1% crystal violet for 1 hour, washed in tap water, dried, and counted. The TCID50/ml were calculated according to published Reed-Muench method [39].

**Table 1.** A summary of the titres of arboviruses in 3 separate titration batches. Each batch represents a biological replicate grown and analysed at a different time point. Titre is expressed as Median Tissue Culture Infectious Dose (TCID50)/ml with the units Virus Particle (VP)/mL.

| Arbovirus | Titre (VP/mL) |  |  |
| --- | --- | --- | --- |
|  | Biological replicate | Biological replicate | Biological replicate |
|  | 1 | 2 | 3 |
| BFV QML | 10<sup>7.75</sup> | 10<sup>7.75</sup> | 10<sup>9.5</sup> |
| BFV WEN | 10<sup>8.3</sup> | 10<sup>7.25</sup> | 10<sup>8.3</sup> |
| RRV QML1 | 10<sup>7.8</sup> | 10<sup>8.25</sup> | 10<sup>8.25</sup> |
| SINV 18953 | 10<sup>8.5</sup> | 10<sup>8.5</sup> | 10<sup>8.5</sup> |
| DENV1 EM-093 | 10<sup>6.75</sup> | 10<sup>6.5</sup> | 10<sup>6.5</sup> |
| DENV2 NGC | 10<sup>7.6</sup> | 10<sup>7.6</sup> | 10<sup>7.6</sup> |
| DENV3 44002 | 10<sup>6.3</sup> | 10<sup>6.3</sup> | 10<sup>6.3</sup> |
| DENV4 AFRIM | 10<sup>6.6</sup> | 10<sup>6.6</sup> | 10<sup>6.6</sup> |

### *P. falciparum* cell culture

Base media prepared for *P. falciparum Welch* was ATCC medium 2196 – Malaria medium (American Type Culture Collection, USA) which consist of a sterilized mixture of RPMI-1640 (Sigma R-0883), HEPES buffer (1 M), Gentamicin (50 mg/ml), L-glutamine (100 mM), Hypoxanthine (100 mM), Glucose (20%) and NaOH (1 N). The mixture was sterilised by filtering through 0.22 μM Millex® filter (Millipore, USA). Complete medium was made by adding the heat-inactivated (at 56°C for 1 h) human plasma to 10% (vol/vol) to the Base medium and was used to culture parasites as previously described [40]. *P. falciparum Welch* had an initial concentration of 255 parasites/mL.

Five biological replicates of serum samples each consisting of 150 mL of pooled human serum samples were obtained from Australian Red Cross Lifeblood using human ethics protocol approved by The University of Queensland (Ethics approval number 2020001077). Following collection from donors, all samples were routinely tested for Hepatitis B and C, HTLV I/II, Syphilis HIV 1/2, and ABO/Rh antigens. Human serum was stored at -25°C for 3-4 days prior to experiment to preserve proteomic profile integrity and was thawed fully at room temperature before use.

### Spectra collection and analysis

LabSpec 4i near-infrared spectrometer (ASD Malvern Panalytical, Malvern, United Kingdom) was used to scan all samples. Details of the spectrometer used is published elsewhere [41]. RS^3^ software (Malvern ASD Panalytical) was used for NIRS spectra collection. Baseline calibration and optimization were done at the beginning of each experiment and thereafter after every 30 minutes by scanning an empty space on the glass slide placed on a white Spectralon plate. Five µL of each arbovirus, *P. falciparum Welch*, and respective media were aliquoted onto glass microscope slides to obtain a sample. A total of 10 technical replicates were scanned for each biological replicate of arbovirus, *P. falciparium* and media. Samples were scanned at approximately 2 mm from the light source by pointing the probe down to the centre of the sample for approximately 3-5 seconds.

### Artificial Neural network analysis

Reflectance spectra were converted to absorbance using the formular 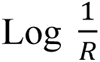. All spectral signatures were converted from txt to csv in ViewSpecPro software (Analytical Spectral Devices Inc, Boulder, CO, USA, 1990–2017). To identify arbovirus and *Plasmodium* peaks of importance, raw spectra was converted into second derivative using the Savitzky–Golay [42] with 2nd order smoothing by combining 10 neighbouring data points. Second derivative spectra graphs were visualised in GraphPad Prism 9 (GraphPad Software, Inc, California, USA, 1989). Model screening and data analysis were conducted in JMP Pro 16 software on raw data (SAS Institute Inc., Cary, NC, USA, 1989–2021). Spectra of DENV1, DENV2, DENV3 and DENV4 were combined into a single identifier referred to as DENV. Likewise, spectra of BFV QML strain and BFV WEN 1631 strain was combined into a single identifier referred to as BFV.

SINDV data was analysed separately due to spectral outliers identified that were causing the model to misclassify other arboviruses. The data was first split into two groups: model training/validation (consisting of 16 biological replicates) and an independent test set (consisting of 8 biological replicates). The training/validation and independent test sets were separate biological replicates grown and analysed at different time points.

**Figure 1.**
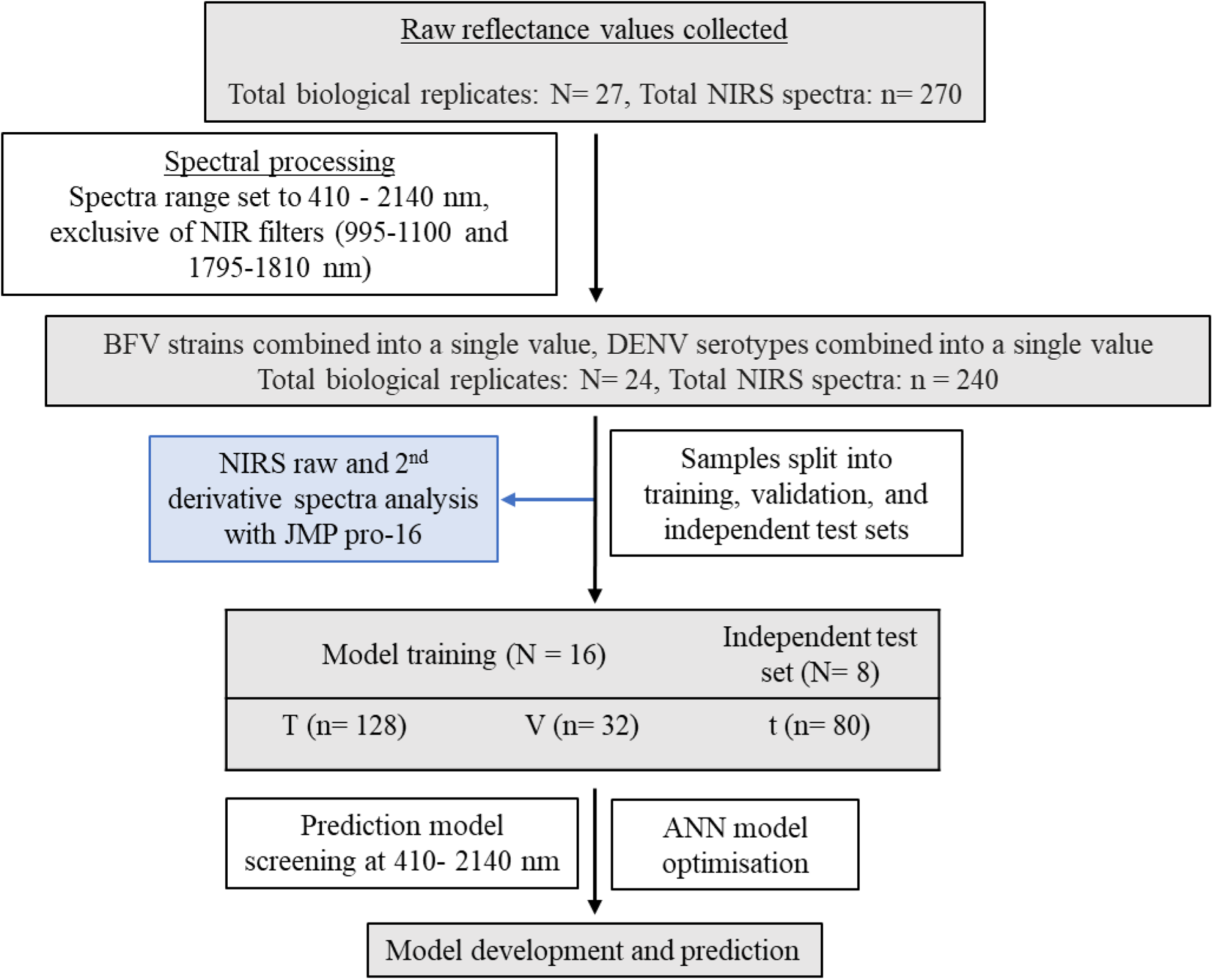
Flow diagram of experimental data. The flow of information from data collection to analysis including the number of samples used for training, validation and test sets. N refers to number of biological replicates and n refers to the number of technical replicates collected and analysed. Training and validation sets were split based on biological replicates. ‘T’ represents training set, ‘V’ represents validation set and t represents the independent test set.

To determine the appropriate machine learning algorithm for each data, raw spectral data underwent model screening where the following model types were screened for preliminary accuracy: Decision Tree [43], Bootstrap Forest [44], Boosted Tree [45], Naïve Bayes [46], Neural Boosted [47] and Support vector machines [48]. Boosted Artificial Neural Network produced the most accurate preliminary results and was therefore selected for further analysis. Spectral signatures from 410 to 2140 nm were used. This region was exclusive of spectral noise between 350-409 and 2141-2500 nm and NIR transition filters at 995-1100 and 1795-1810 nm were used as model predictors whereas infection (arbovirus) was used as the response factor. ANN model was developed using random K-Fold cross-validation (n=5 samples). The Neural Networks consisted of one layer with three TanH activation nodes boosted at a learning rate of 0.1 iteratively for 100 tours. Models were built to differentiate arboviruses from media. A summary of sample distribution between training, validation and independent test set is shown in Figure 1.

## Results

### NIRS arboviruses 2^nd^ derivative spectra

Signature peaks for DENV are as seen at 410 nm within the visible region and at 1130 nm within the NIR region. DENV peak is observed to have lower absorbance than media at 410 nm but a higher absorbance value than media at 1130 nm (Fig 2A).

A total of 10 prominent peaks were identified for BFV. These prominent peaks were seen within the visible region at 410, 430, 562 and 588 nm and within the NIR region at 946, 958, 1130, 1154, 1287-1331 and 1780 nm. Absorbance peaks within the visible region at 430 and 588 nm and within the NIR region at 946, 1130 nm were seen to have higher absorbance values than media while peaks at 410 and 562 nm in the visible region and 958, 1154 and 1780 nm in the NIR region were observed to have lower absorbance values than media (Fig 2B).

For RRV, prominent peaks were identified at 410 and 430 within the visible region and at 1130, 1154, 1447, 1464 and 1780 nm within the NIR region. Overall, RRV was observed to have 7 prominent peaks (Fig 2C).

The second derivative NIR spectra of SINV 18953 showed prominent peaks at 412, 1447 and 1463 nm. SINV had the lowest number of virus specific prominent peaks (3 peaks) compared to the other arboviruses. Unique virus peaks were observed at 410,1447 and 1463 nm (Fig 2D).

**Figure 2.**
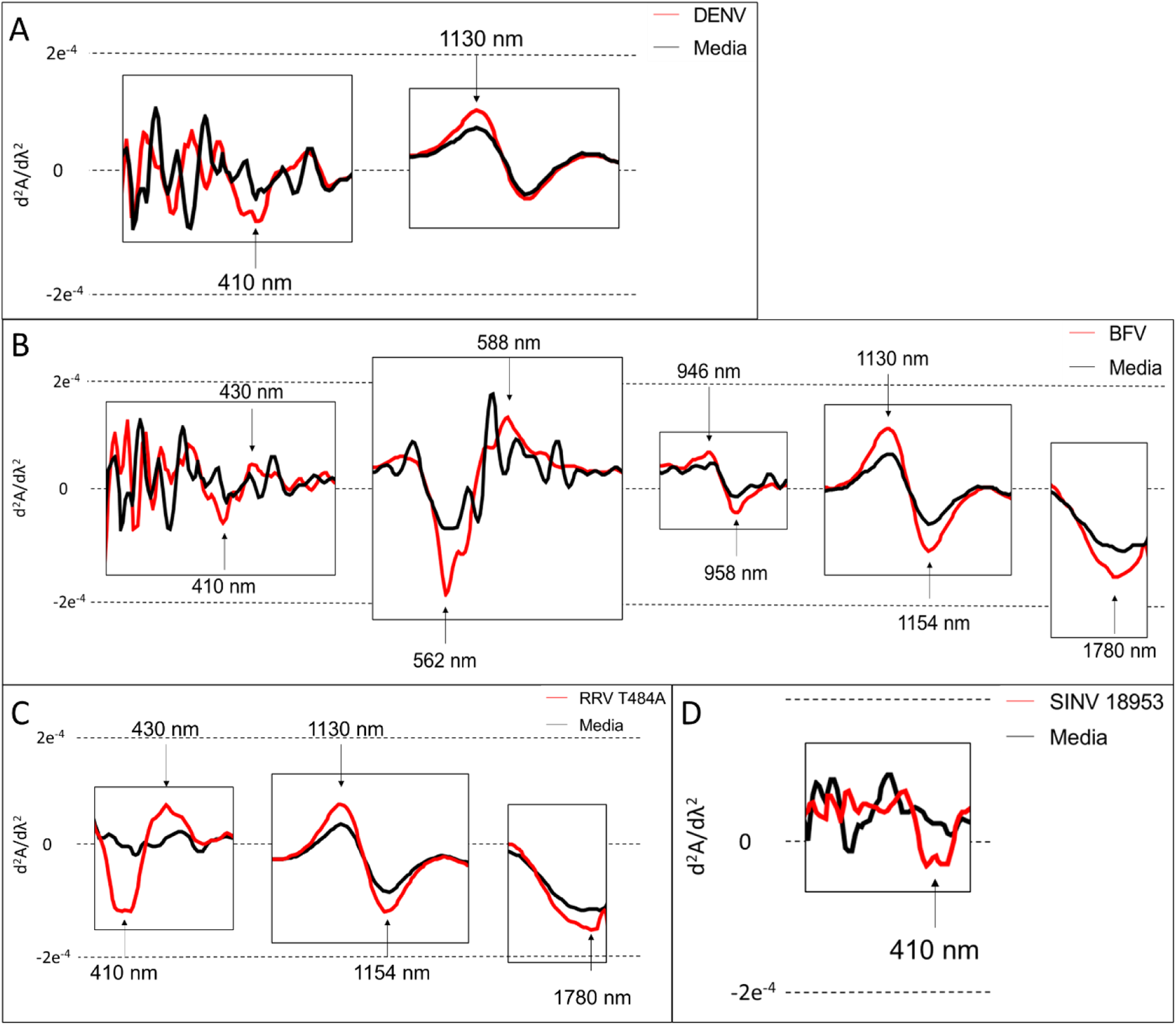
The average 2^nd^ derivative of visible and NIR spectra highlighting prominent peaks of DENV and media (A), BFV/media (B), RRV/media (C) and SINV/media (D). Peaks of importance are shown with black arrows.

### Summary of DENV, BFV, SINV and RRV signature NIR peaks

Four pathogen related wavelengths fell within the visible light region. Half of which (410 and 430 nm) fell within the blue visible region while the rest (562, 588 nm) fell in the green visible light spectrum. BFV prominent peaks at 946 and 958 nm was observed in the 3^rd^ overtone region. Three pathogen related prominent peaks (1130, 1154 and 1780 nm) was seen in the 2^nd^ overtone region. Peaks at 410, 430, 1130 and 1780 nm was observed in more than one arbovirus. A summary of all arboviruses with their proposed functional group/ vibrations and molecular structure relative to published literature is shown in Table 2.

**Table 2.** Summary of unique 2^nd^ derivative arbovirus peaks, their respective functional group or vibration and representation from literature [49–54].

| Wavelength (nm) | Arbovirus | Wavelength region | Functional group/<br>vibration |
| --- | --- | --- | --- |
| 410 | BFV strains, DENV strains, RRV and SINV | Visible light | Blue visible light |
| 430 | BFV strains and RRV |  |  |
| 562 | BFV strains |  | Green visible light |
| 588 |  |  |  |
| 946 | BFV strains | 3<sup>rd</sup> overtone region | CH<sub>2</sub> |
| 958 |  |  |  |
| 1130 | BFV strains, DENV strains and RRV | 2<sup>nd</sup> overtone region | CH<sub>3</sub> |
| 1154 | BFV strains |  |  |
| 1780 | BFV strains and RRV |  | C=O |

### ANN differentiation of BFV, DENV, RRV, SINV and Vero media

To identify if machine learning could differentiate arboviruses from Vero media samples using spectra collected, a model using ANN was applied on the raw spectra collected i.e., 410-2140 nm (Table 3). Overall, the ANN differentiated BFV, DENV, RRV and Vero media from each other with an. R square value of 1 for both training (n= 128) and validation (n= 32) set (Table 3). The independent test set consisting of 80 NIR spectra was predicted using the training model. Overall, positive predictive rate defined as the ratio of samples truly predicted as positive out of those that were predicted as positive was 100% indicating all infected samples were predicted as infected. The negative predictive rate defined as the samples predicted as negative out of all those that are negative was 76.9% (n= 13) meaning some positive samples were predicted as not infected. Specificity defined as the proportion of samples predicted as negative out of all negative samples was 100% (n= 10) meaning all media samples were predicted as not infected. Summary of the training, validation and independent test set is shown in table 3.

**Table 3.** A confusion matrix showing the accuracy of differentiating arboviruses from each other and from media for the training, validation, and independent test sets.

| Training set |  | Prediction count |  |  |
| --- | --- | --- | --- | --- |
| Actual | BFV | RRV | DENV | Media |
| BFV | 32 | 0 | 0 | 0 |
| RRV | 0 | 16 | 0 | 0 |
| DENV | 0 | 0 | 64 | 0 |
| Media | 0 | 0 | 0 | 16 |
| Validation set |  | Prediction count |  |  |
| Actual | BFV | RRV | DENV | Media |
| BFV | 8 | 0 | 0 | 0 |
| RRV | 0 | 4 | 0 | 0 |
| DENV | 0 | 0 | 16 | 0 |
| Media | 0 | 0 | 0 | 4 |
| Independent test set |  | Prediction count |  |  |
| Actual | BFV | RRV | DENV | Media |
| BFV | 14 | 1 | 3 | 2 |
| RRV | 0 | 10 | 0 | 0 |
| DENV | 7 | 0 | 32 | 1 |
| Media | 0 | 0 | 0 | 10 |

**Table 4.**
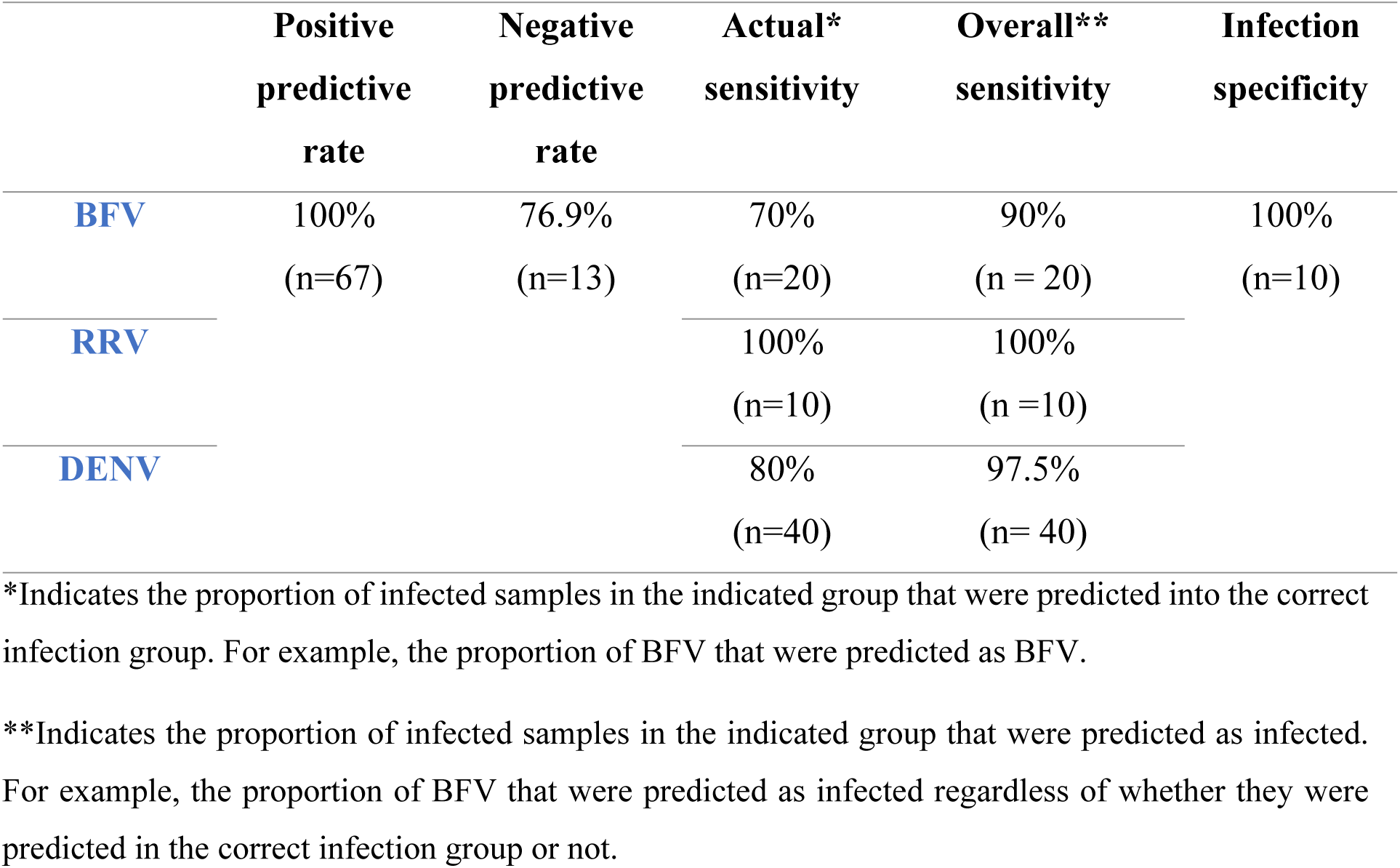
A summary of positive prediction rate, negative prediction rate, sensitivity, and specificity of BFV, RRV and DENV. Arbovirus Type.

|  | Positive<br>predictive<br>rate | Negative<br>predictive<br>rate | Actual*<br>sensitivity | Overall**<br>sensitivity | Infection<br>specificity |
| --- | --- | --- | --- | --- | --- |
| BFV | 100%<br>(n=67) | 76.9%<br>(n=13) | 70%<br>(n=20) | 90%<br>(n = 20) | 100%<br>(n=10) |
| RRV |  |  | 100%<br>(n=10) | 100%<br>(n =10) |  |
| DENV |  |  | 80%<br>(n=40) | 97.5%<br>(n= 40) |  |

\*Indicates the proportion of infected samples in the indicated group that were predicted into the correct infection group. For example, the proportion of BFV that were predicted as BFV.
\*\*Indicates the proportion of infected samples in the indicated group that were predicted as infected. For example, the proportion of BFV that were predicted as infected regardless of whether they were predicted in the correct infection group or not.

### NIRS raw spectra for *P. falciparum*

Unique NIR peaks of *P. falciparum* were observed in the 350-650nm visible region and 1450, 1960 nm NIR region (Fig 3A). Relative to media, a unique peak for *P. falciparum* was observed at 440 and 543 nm*. P. falciparum* also showed a higher absorbance than media at both 1450 and 1960 nm. To further investigate the spectra, we plotted the 2^nd^ derivative of the average spectra. 2^nd^ derivative spectra of *P. falciparum*. Peaks related to *P. falciparum* were seen at 514, 528, 547, 561, 582 and 595 within the visible region 1388, 1432, 1681, 1700, 1721, 1882, 1905, 2245, 2278 and 2300 nm within the NIR region (Fig 3B). Some of the wavelengths observed to be unique for *P. falciparum* are found within the visible light region at 514, 528, 547, 561, 582 and 595 nm. The highest peaks for *P. falciparum* in this region were at 582 and 595 nm. (Fig 3B). Comparatively, no peaks were observed for media used to grow *P. falciparum* in the visible region.

**Figure 3.**
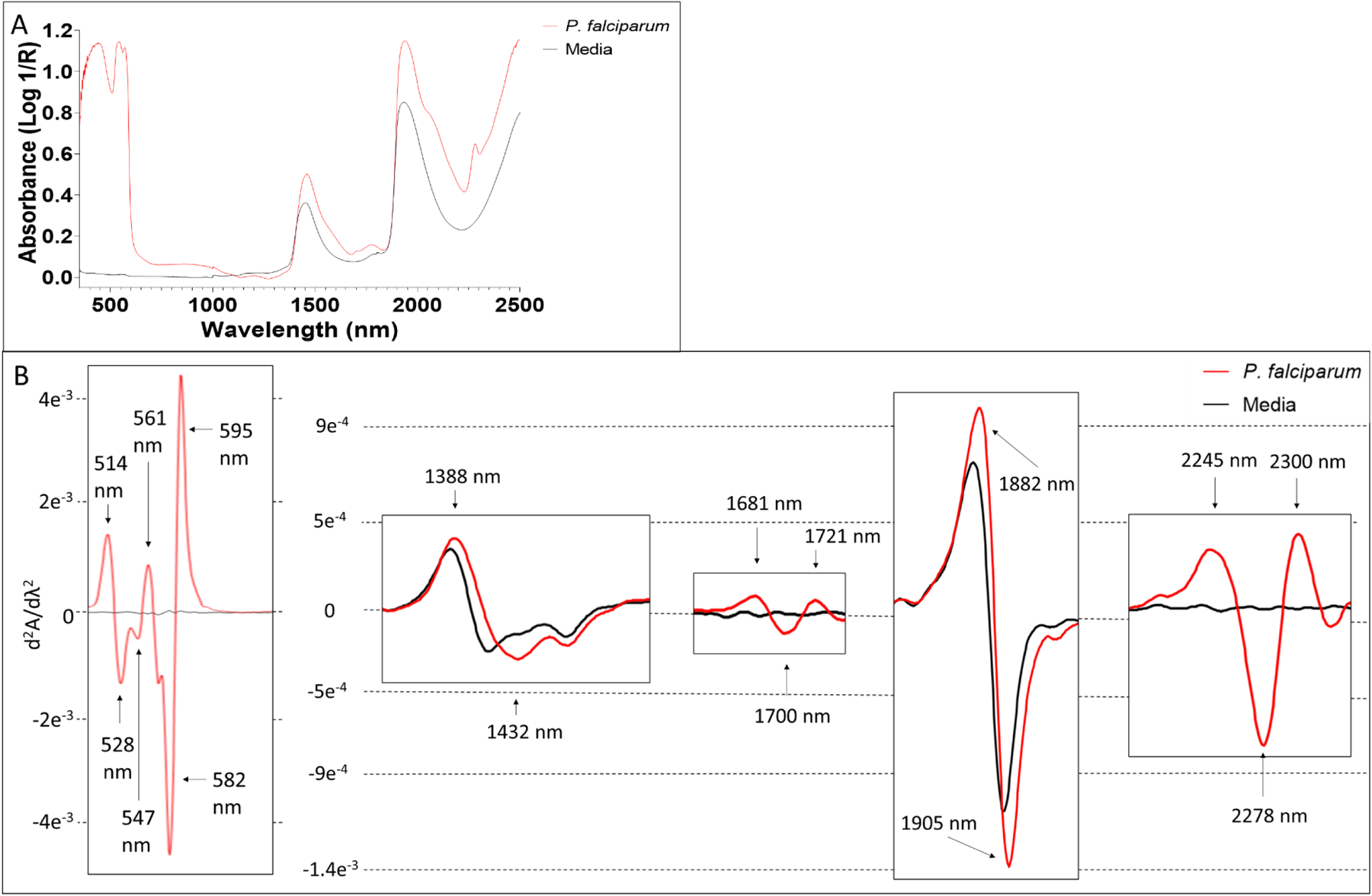
The average raw visible NIR spectra (A) and the second derivative spectra (B) of *P. falciparum* and media. Peaks of importance are indicated with black arrows.

### Summary of *P. falciparum* signature NIR peaks

Six peaks observed for *P. falciparum* belong to the visible light region, two within the 2^nd^ overtone region, five in the 1^st^ overtone region and 3 in the combination band region. The majority of the peaks identified belong to the visible light region and the 1^st^ overtone region (Table 5).

**Table 5.**
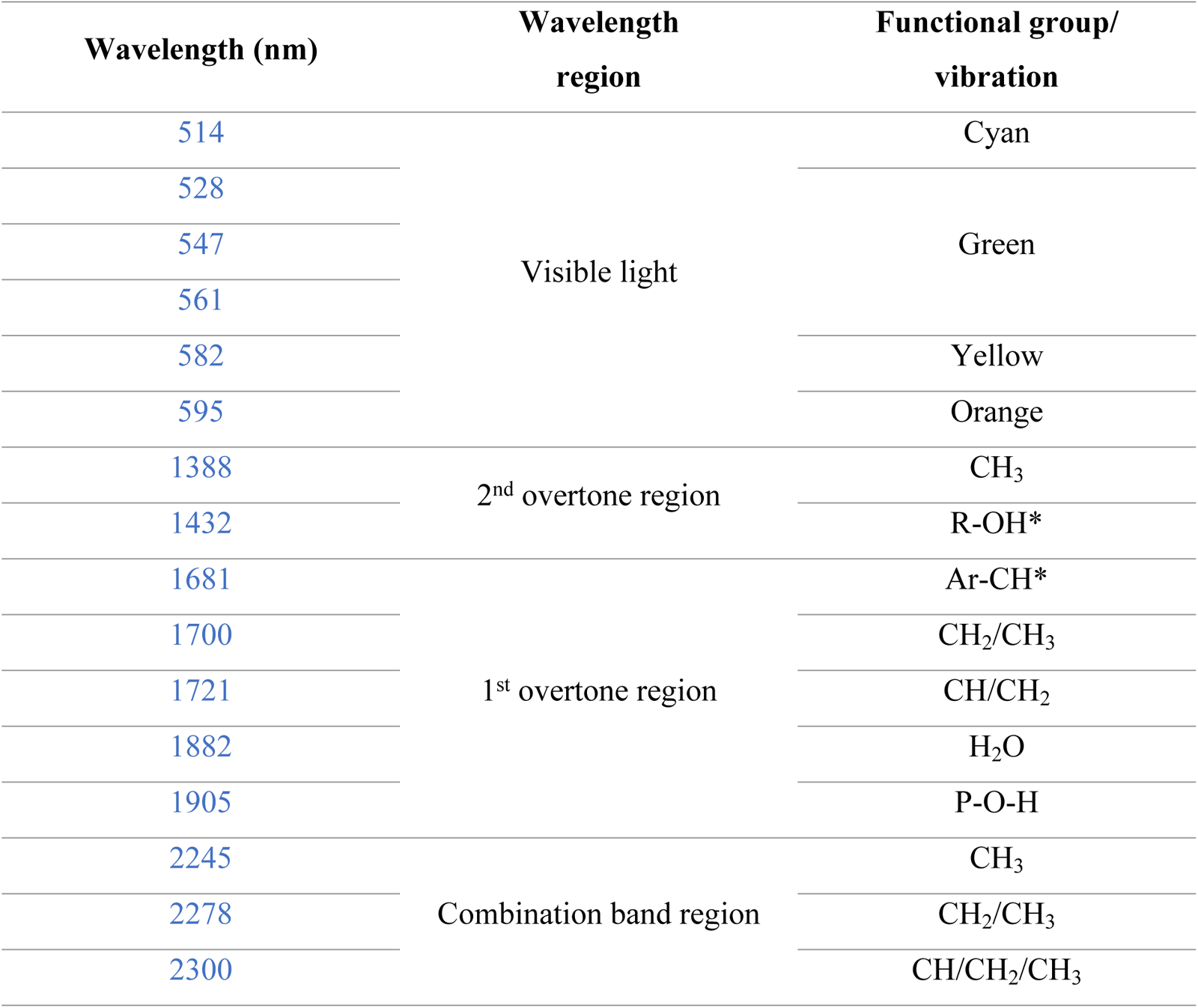
Summary of peaks identified for *P. falciparum* their respective functional group relative to literature [49–54].

## Discussion

The aim of this study was to identify visible and NIR peaks of importance for some arboviruses and *P. falciparum* as potential diagnostic biomarkers for these infections. A total of 20 peaks of interest were identified for the arboviruses tested in this study (Table 3). Distinct signature peaks for BFV were seen at 562, 588, 946, 958 and 1154 nm. Peaks at 562 and 588 nm are within the visible region and could be useful for identification of BFV. Peaks at 946, 958 and 1154 nm represent lipid molecular structures. The lipids identified could be due to the presences of a bilipid membrane anchored on the surface of BFV by proteins E1 and E2 [55].

Besides BFV, the rest of the unique peaks observed were all shared among the three arboviruses and the peak at 410 nm was observed in all arboviruses. A peak at 410 nm is commonly used for assays that require fluorescent excitation such as ELISA and microscopy [56–59]. DENV and RRV have been identified using this peak previously using a spectrofluorometer [60] and ELISA [61], respectively. A peak at 430 nm was observed for BFV strains and RRV but not DENV and SINV indicating the likelihood that visible light could possibly be used to distinguish between these arboviruses. Peaks at 1130 and 1780 nm were present in BFV strains and RRV. The peak at 1130 nm represents CH_3_ functional group whereas the peak at 1780 nm represents the C=O functional group. Both wavelengths represent the presence of lipid molecules [49–54]. For BFV, this peak could be due to the presence of the bilipid membrane [55]. RRV uses lipid droplet biogenesis for viral replication [62] and lipid rafts for infection [63] thus the lipids identified here could be the residue from these processes. The peak at 1130 nm was also observed for DENV. Lipids present in DENV could be a by-product from cellular passaging of DENV in C6/36 *Ae. albopictus* cells which use lipid metabolism for efficient replication [64–66]. In addition, we have identified this peak in a previous study to detect DENV1 in human blood plasma [67].

To further evaluate if the NIRS spectra could differentiate between arboviruses and media, machine learning algorithms were run on the visible-NIR spectral signatures collected. SINV spectral data was excluded from the training model because only one unique peak (410 nm) was identified, and this peak was not unique to SINV causing misclassification. Using the ANN model, the sensitivity and specificity of ≥90% and 100%, respectively were achieved for the independent test set when samples were grouped as infected or not infected. However, sensitivity for predicting the arboviruses into their actual group was 100%, 80% and 70% for RRV, DENV and BFV respectively. Seven out of 40 DENV samples were predicted as BFV. Similarly, 3 out of 20 BFV samples were predicted as DENV (Table 3). This indicates a slight confusion in the model between DENV and BFV which could be due to the shared peaks at 410 and 1130 nm.

A total of 16 peaks were identified for *P. falciparum* six of which belong to the visible light region. This is not surprising as the parasite can be detected via light microscopy. Three of the 6 peaks within the visible light region; 547, 561 and 582 nm are within the same region as peaks identified in the ring stage of *P. falciparum* in whole blood as reported by another study (540, 560 and 579 nm) [27]. Of the 10 remaining wavelengths, 3 in the NIR region (1388, 2245 and 2300 nm) represent C-H bond vibrations [49–54]. C-H bonds are basic chemical building blocks of life and could be responsible for numerous structures within the *P. falciparum* parasite. Four wavelengths of interest represent lipids [49–54]. Lipids have been found in *P. falciparum* and have been shown to play several roles such as toxicity activity [68], gametocytogenesis [69], and development [70–73]. In addition, peaks at 1388 and 1432 nm are within the same range as those recently identified (1377 and 1431 nm) identified in malaria infected patients non-invasively using a handheld spectrometer [74].

The prominent peaks for arboviruses and *P. falciparium* detected in this study will be useful for developing NIR library of peaks for disease surveillance/diagnosis. If further assessed these arboviruses and *P. falciparium* could be developed into biomarkers of these pathogens in human samples. NIRS is an easy to use, rapid, potable, and reagent free technique [67, 75–77]. Once machine learning techniques are developed, the feasibility for on-site largescale diagnosis of patients is feasible. With this additional tool in hand, NIRS has the potential to rapidly identify infections to stop an outbreak by facilitating timely isolation and treatment of patients.

## Conclusion

In this study we have identified several unique and novel visible and NIR peaks for BFV, DENV, RRV, SINV and *P. falciparum*. We have also demonstrated the ability of NIRS and machine learning to differentiate between arboviruses from their media. To our knowledge this is the first investigation to report arbovirus NIR peaks that have potential for arbovirus identification. The findings of this study are important as they provide insights into the potential future application of these peaks as diagnostic biomarkers for these pathogens. Future work should evaluate the capability of NIRS to detect these pathogens using these biomarkers in human blood samples.

